# Synthetic autotrophic yeast enables high itaconic acid production from CO_2_ via integrated pathway and process design

**DOI:** 10.1101/2025.06.16.659842

**Authors:** Özge Ata, Lisa Lutz, Michael Baumschabl, Diethard Mattanovich

**Affiliations:** BOKU University, Vienna, Department of Biotechnology and Food Science, Institute of Microbiology and Microbial Biotechnology, 1190 Vienna, Austria; Austrian Centre of Industrial Biotechnology, Vienna, 1190, Austria

## Abstract

Single carbon (C1) substrates are gaining importance as future feedstocks for the production of bio-based chemicals. Carbon dioxide, a major greenhouse gas, offers a promising alternative to the traditional feedstocks to shift towards C1-based, sustainable processes. Here, we present a synthetic autotrophic *Komagataella phaffii* (*Pichia pastoris*) that is able to produce itaconic acid by the direct conversion of CO_2_, achieving final titers of approximately 12 g L^-1^ in bioreactor cultivations. We show that a combined approach that integrates balancing the flux between the Calvin–Benson–Bassham (CBB) cycle and itaconic acid metabolism with process design was essential to enhance the production. Our study demonstrates the potential of *K. phaffii* as a microbial platform using CO_2_ as the direct carbon source, aligning with the future goals of establishing sustainable bioprocesses.

## Introduction

As we struggle with the challenges of the climate crisis, it is essential to develop innovative strategies that not only mitigate these issues but also offer solutions for long-term sustainability. Carbon dioxide (CO_2_), one of the major greenhouse gases, is increasing in atmospheric concentration due to societal activities, which endangers our entire planet. In this regard, single carbon (C1)-based bioprocesses emerge as a promising technology to reduce CO_2_ emissions, via the microbial conversion of CO_2_ or other CO_2_-derived C1-sources such as methanol and formate.^1–3^

Several natural or engineered autotrophs^4–8^ and acetogens^9–12^ have been harnessed to convert CO_2_ into a broad range of industrially relevant chemicals such as alcohols and organic acids.^13–17^ Among these organic acids, itaconic acid has gained interest worldwide as being one of the 12 top-value added chemicals by the US Department of Energy. The global market size of production of itaconic acid has reached more than 100 million USD in 2024 and expected to exceed more than 170 million USD by 2031.^18^ Itaconic acid can serve as a building block for several products such as plastics, drug carriers, polymer binding agents, resins, and synthetic fibers.^19^ Conventionally, *Aspergillus terreus* is the native, predominant industrial host for the microbial production of itaconic acid.^20,21^ The biosynthesis pathway of itaconic acid involves the key gene cis-aconitate decarboxylase, *cadA*, which converts *cis*-aconitate into itaconic acid (Figure 1). Two additional transporters, mitochondrial *cis*-aconitate transporter (MttA) and the major facilitator superfamily transporter (MfsA), transports the substrate cis-acotinate from the mitochondria to the cytosol, or the product itself from the cytosol to the extracellular environment, respectively. However, the industrial microbial production of itaconic acid by *A. terreus* predominantly relies on sugar-based feedstocks, competing with agricultural land use. A C1-substrate based bioprocess could therefore play a key role in enabling more sustainable production.

**Figure 1.**
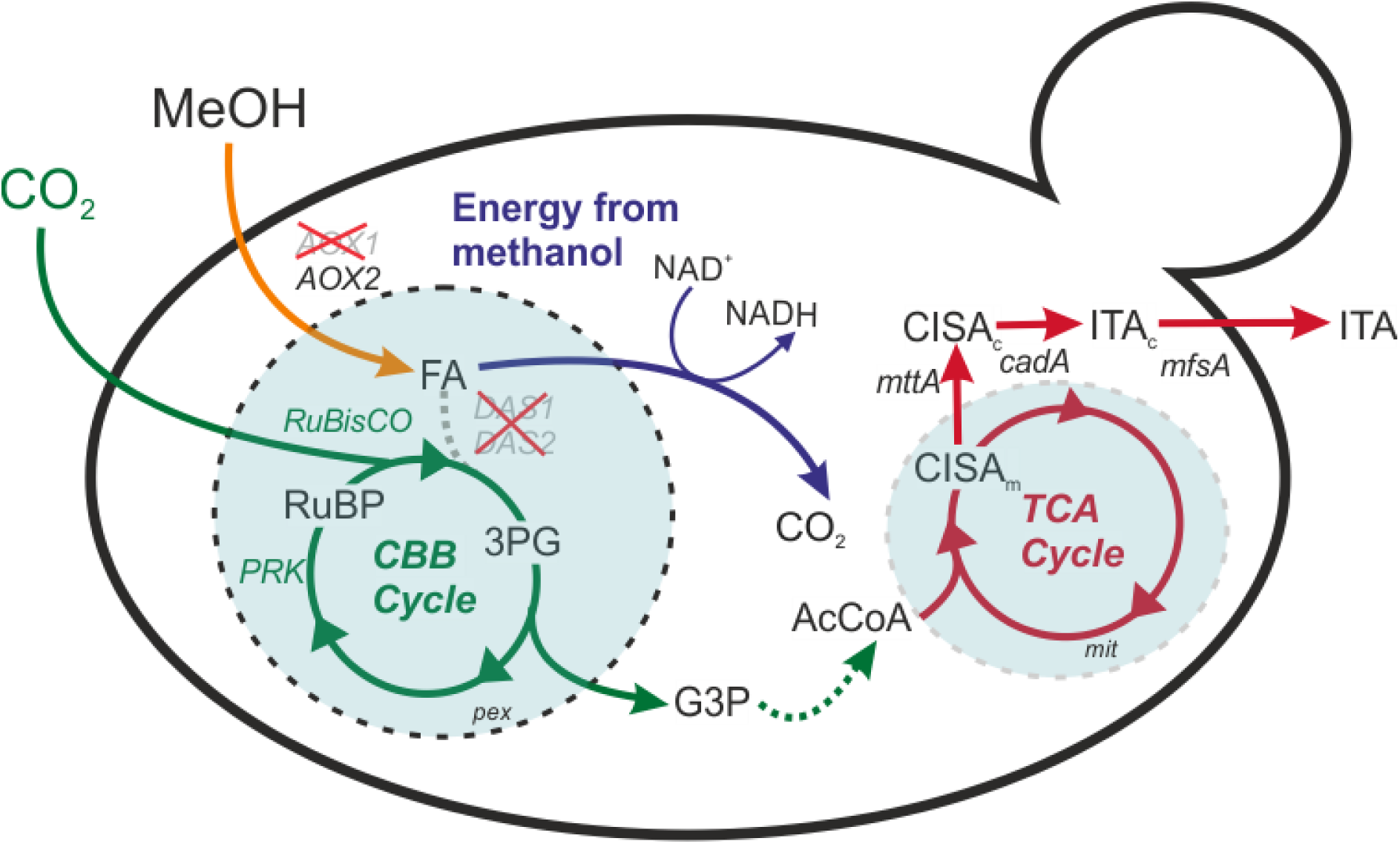
Metabolic pathway of the itaconic acid production in the synthetic autotrophic *K. phaffii.* Deletion of *DAS1* and *DAS2* interrupts methanol assimilation (dashed gray line). *AOX1* was knocked out to reduce the rate of formaldehyde formation which could be toxic to the cells. More details about the engineering strategy can be found in 22. Expression of the key enzyme *cadA* among with two transporters, *mttA* and *mfsA*, enables itaconic acid production from CO_2_. 3PG: 3-phosphoglycerate, AcCoA: acetyl-coenzyme A, *AOX1* and *AOX2*: alcohol oxidase 1 and 2, *cadA*: cis-aconitate decarboxylase, CBB cycle: Calvin-Benson-Bassham cycle, CISA_c_: cytosolic cis- aconitate, CISA_m_: mitochondrial cis-aconitate, *DAS1* and *DAS2*: dihydroxyacetone synthase 1 and 2, FA: formaldehyde, G3P: glyceraldehyde 3-phosphate, ITA: itaconic acid, *mttA*: mitochondrial tricarboxylic acid transporter, NAD+/NADH: nicotinamide adenine dinucleotide, *PRK*: phosphoribulokinase, RuBP: ribulose 1,5-bisphosphate, *RuBisCO*: ribulose 1,5-bisphosphate carboxylase/oxygenase. pex: peroxisome, mit: mitochondria.

In a recent study, we demonstrated that a synthetic autotrophic yeast *Komagataella phaffii* (*Pichia pastoris*), can produce itaconic acid up to 2 g L^−1^ titers in shake flask cultivations using CO_2_ and methanol as the sole carbon and energy sources, respectively; through the introduced CBB cycle. However, upscaling proved challenging, as titers in lab-scale bioreactors reached only around 0.5 g L⁻¹, underlining the need for rigorous process design to achieve higher production levels.

In the present study, we seek to increase the itaconic acid production performance of the synthetic autotrophic *K. phaffii* from CO_2_ in lab-scale bioreactor cultivations. Through combined efforts of metabolic engineering to balance the flux of the CBB cycle and itaconic acid metabolism with process parameter optimization, we could achieve 11.84 g L^−1^ ± 0.26 of itaconic acid improving the final titer and specific productivity by 22.5-fold and 5.3-fold respectively, compared to the previous bioreactor cultivation.^16^

## Results

### Dissolved oxygen concentration is one of the key parameters

Recently, we have demonstrated that the production of itaconic acid is achievable through the conversion of CO_2_ by an engineered autotrophic *K. phaffii* strain.^16^ *K. phaffii* is not able to produce itaconic acid as it does not possess a *cadA* gene. Itaconic acid production was enabled by the expression and balancing of *cadA* and *mttA*, resulting in the cadA+mttA strain with a final titer of approximately 0.80 g L^-1^ in shake flasks. Furthermore, it was demonstrated that dissolved oxygen (DO) concentration is a crucial parameter in laboratory-scale fermentations and shown that at high DO concentration (20%), itaconic acid production is lower. Therefore, in the present study, we investigated the setpoints 4%, 8%, and 16% (Figure 2). The growth profile exhibited a uniform pattern across all conditions in bioreactors, with no observable growth, despite the absence of any growth impairments in the tested strain during the shake flask cultivations. However, an obvious trend was observed in the itaconic acid production profiles across varying dissolved oxygen concentrations. The highest titers (0.91 g L^-1^) were observed at 16% O_2_, while the lowest was recorded at 4% (0.65 g L^-1^). Furthermore, analysis of the expression levels of key genes involved in the CBB cycle and itaconic acid metabolism revealed no significant trend in response to varying DO values (Supplementary Figure 1). Consequently, 16% DO was selected for subsequent bioreactor experiments, as this condition yielded the highest itaconic acid titers and productivity (Table 1).

**Figure 2.**
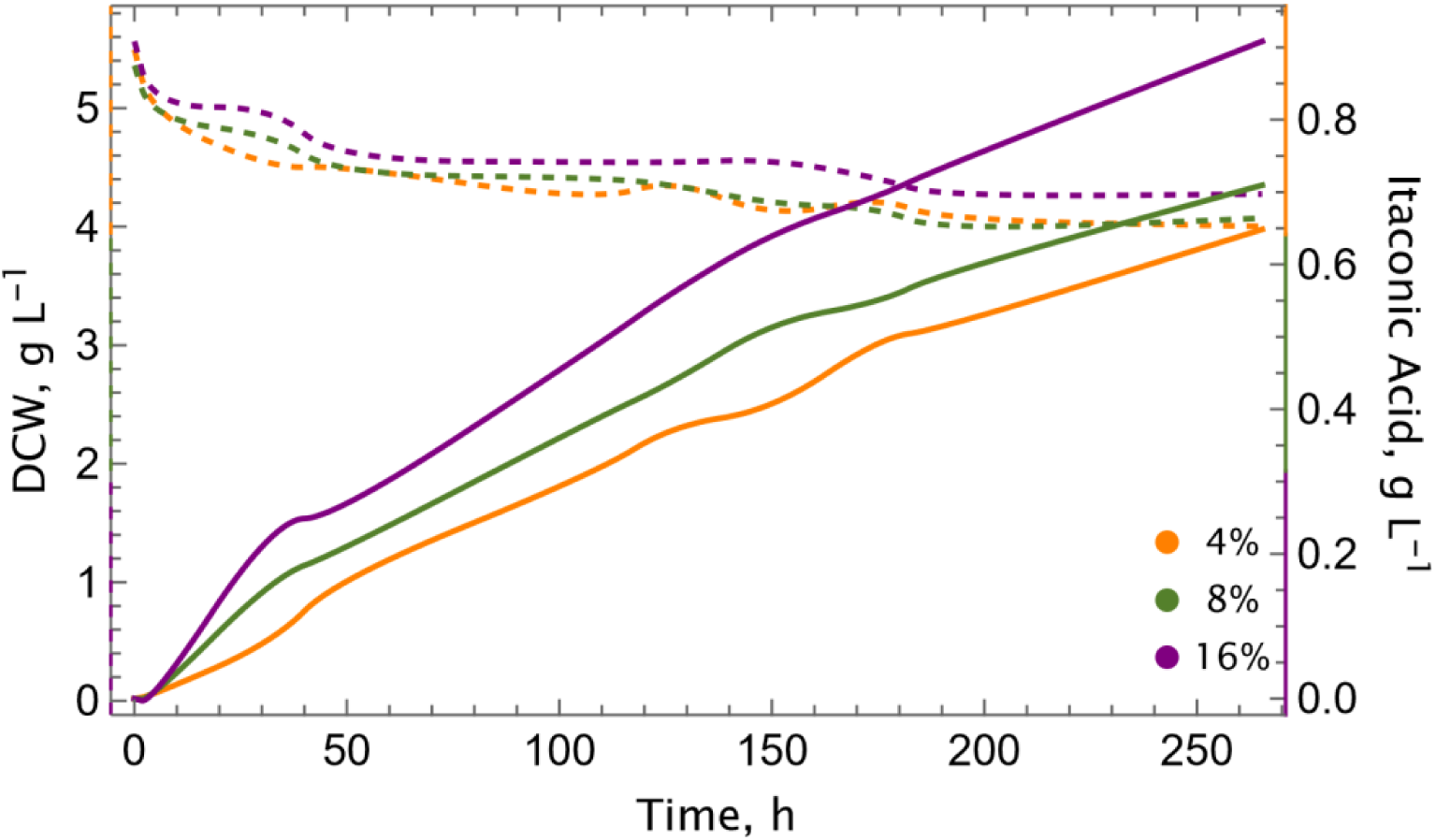
Dissolved oxygen concentration is a key parameter for the production of itaconic acid. Growth (dashed lines) and itaconic acid production (solid lines) profiles of the cadA+mttA strain at different dissolved oxygen concentrations. Experiments were conducted in single runs in lab-scale bioreactors, using 5% CO_2_ in the inlet gas stream, at 30°C.

**Table 1.**
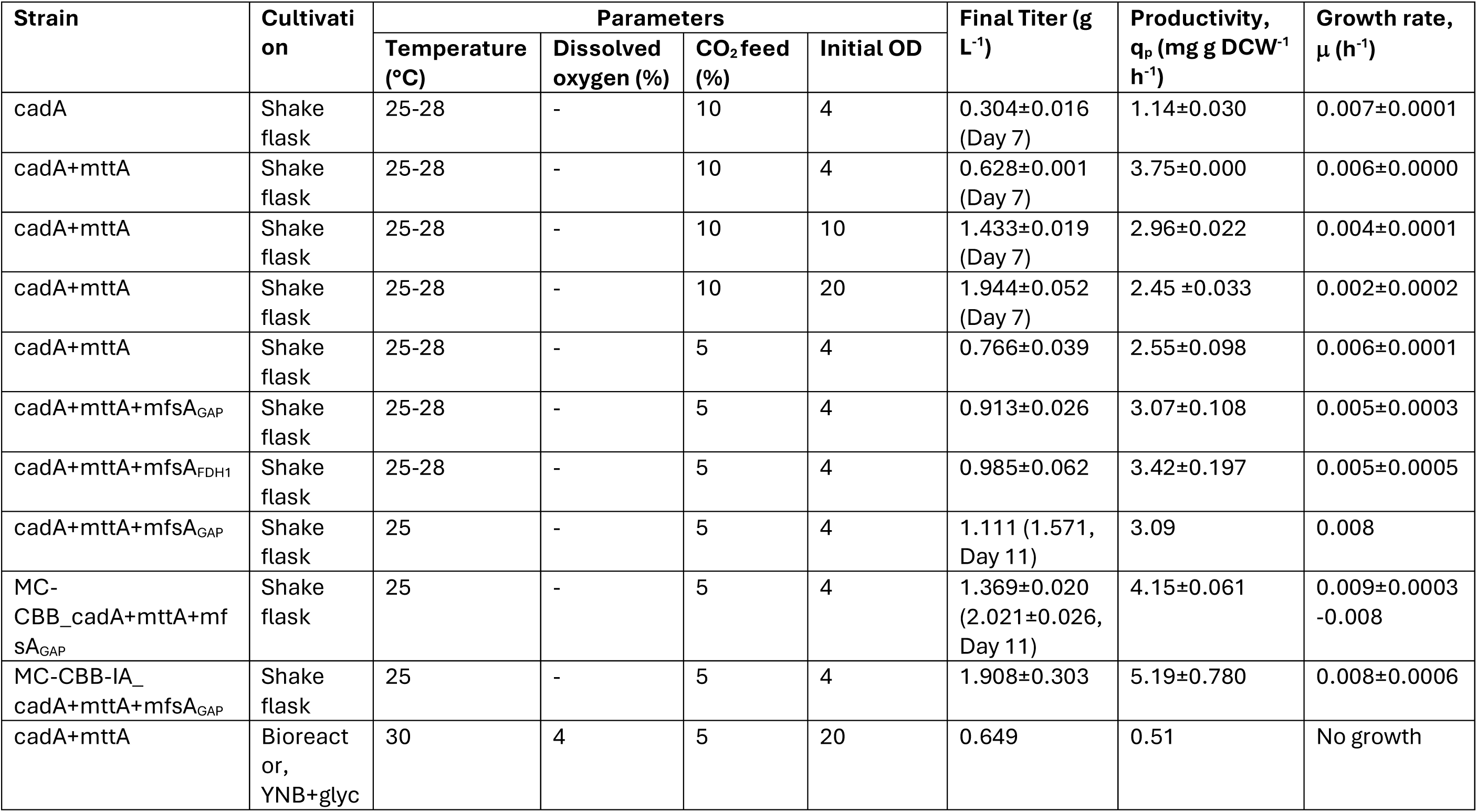
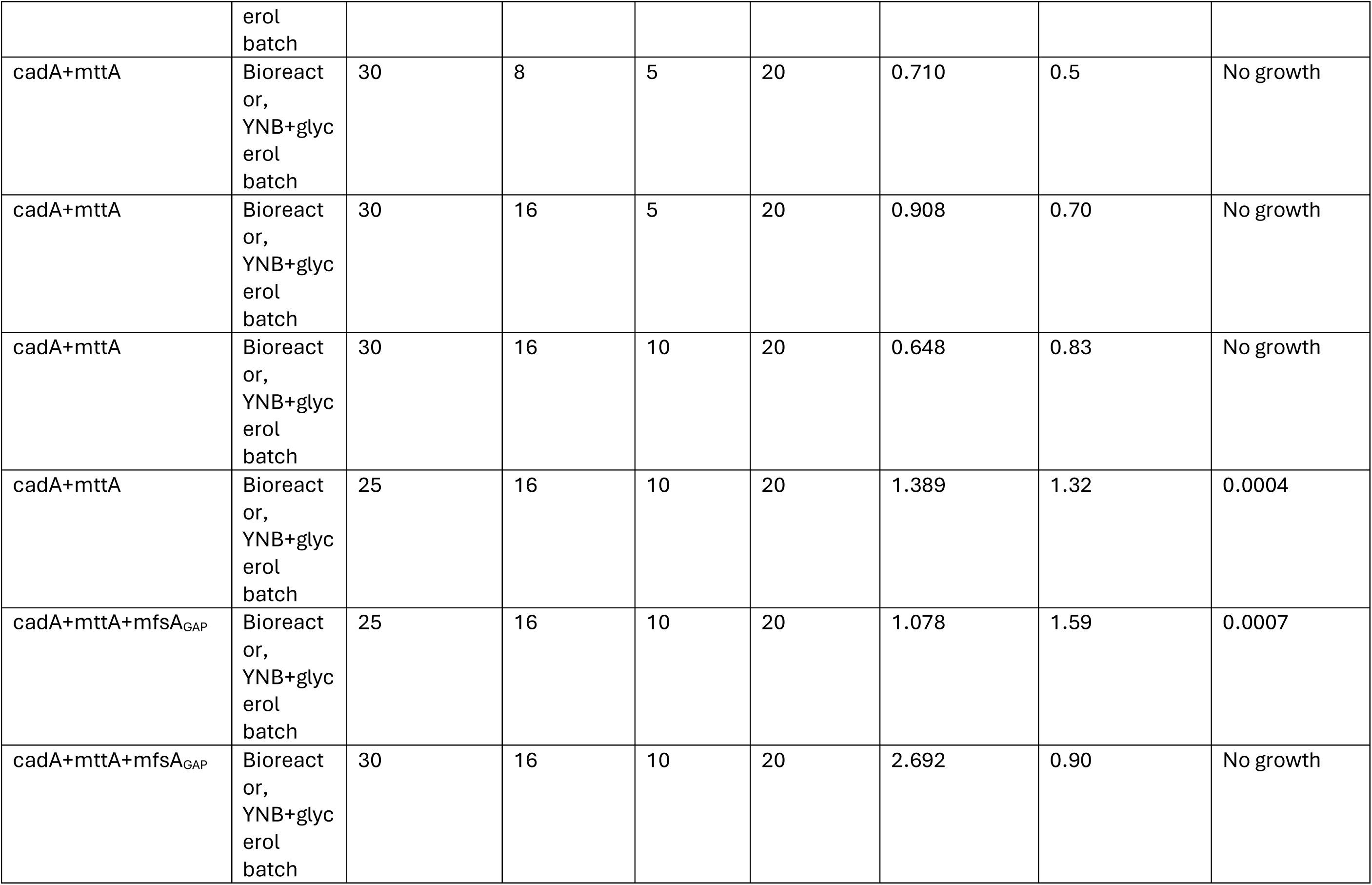
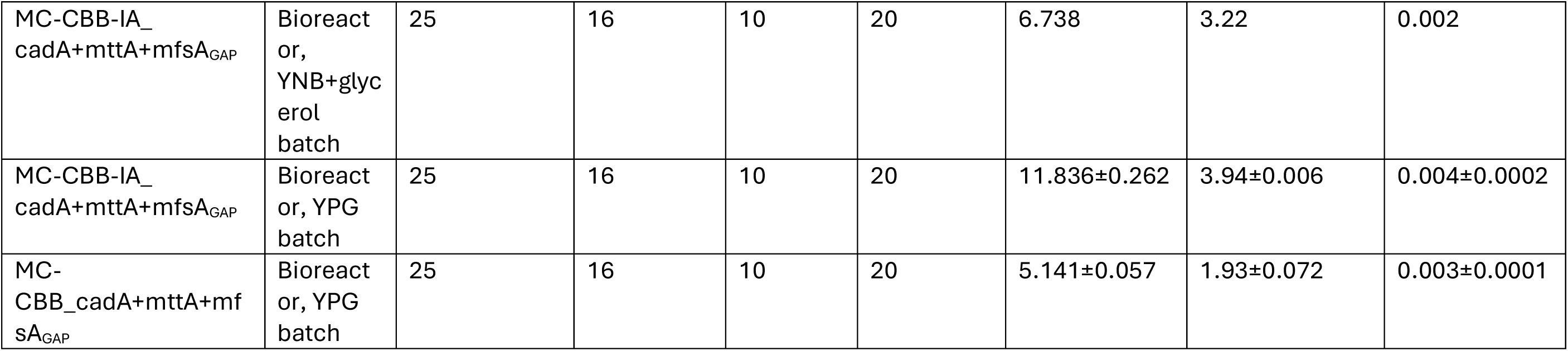
Process parameters in shake flask and bioreactor cultivations. Unless indicated in parantheses, final titers of Day 8 of the cultivation are given. Dissolved oxygen concentration is not controlled during shake flask cultivations. MC: multicopy

| Strain | Cultivation | Parameters |  |  |  | Final Titer (g L<sup>-1</sup>) | Productivity, q<sub>p</sub> (mg g DCW<sup>-1</sup> h<sup>-1</sup>) | Growth rate, μ (h<sup>-1</sup>) |
| --- | --- | --- | --- | --- | --- | --- | --- | --- |
|  |  | Temperature (°C) | Dissolved oxygen (%) | CO<sub>2</sub> feed (%) | Initial OD |  |  |  |
| cadA | Shake flask | 25-28 | - | 10 | 4 | 0.304±0.016 (Day 7) | 1.14±0.030 | 0.007±0.0001 |
| cadA+mttA | Shake flask | 25-28 | - | 10 | 4 | 0.628±0.001 (Day 7) | 3.75±0.000 | 0.006±0.0000 |
| cadA+mttA | Shake flask | 25-28 | - | 10 | 10 | 1.433±0.019 (Day 7) | 2.96±0.022 | 0.004±0.0001 |
| cadA+mttA | Shake flask | 25-28 | - | 10 | 20 | 1.944±0.052 (Day 7) | 2.45 ±0.033 | 0.002±0.0002 |
| cadA+mttA | Shake flask | 25-28 | - | 5 | 4 | 0.766±0.039 | 2.55±0.098 | 0.006±0.0001 |
| cadA+mttA+mfsA<sub>GAP</sub> | Shake flask | 25-28 | - | 5 | 4 | 0.913±0.026 | 3.07±0.108 | 0.005±0.0003 |
| cadA+mttA+mfsA<sub>FDH1</sub> | Shake flask | 25-28 | - | 5 | 4 | 0.985±0.062 | 3.42±0.197 | 0.005±0.0005 |
| cadA+mttA+mfsA<sub>GAP</sub> | Shake flask | 25 | - | 5 | 4 | 1.111 (1.571, Day 11) | 3.09 | 0.008 |
| MC-CBB_cadA+mttA+mfsA<sub>GAP</sub> | Shake flask | 25 | - | 5 | 4 | 1.369±0.020 (2.021±0.026, Day 11) | 4.15±0.061 | 0.009±0.0003 -0.008 |
| MC-CBB-IA_cadA+mttA+mfsA<sub>GAP</sub> | Shake flask | 25 | - | 5 | 4 | 1.908±0.303 | 5.19±0.780 | 0.008±0.0006 |
| cadA+mttA | Bioreactor, YNB+glyc erol<br>batch | 30 | 4 | 5 | 20 | 0.649 | 0.51 | No growth |
| cadA+mttA | Bioreact<br>or,<br>YNB+glyc<br>erol<br>batch | 30 | 8 | 5 | 20 | 0.710 | 0.5 | No growth |
| cadA+mttA | Bioreact<br>or,<br>YNB+glyc<br>erol<br>batch | 30 | 16 | 5 | 20 | 0.908 | 0.70 | No growth |
| cadA+mttA | Bioreact<br>or,<br>YNB+glyc<br>erol<br>batch | 30 | 16 | 10 | 20 | 0.648 | 0.83 | No growth |
| cadA+mttA | Bioreact<br>or,<br>YNB+glyc<br>erol<br>batch | 25 | 16 | 10 | 20 | 1.389 | 1.32 | 0.0004 |
| cadA+mttA+mfsA<sub>GAP</sub> | Bioreact<br>or,<br>YNB+glyc<br>erol<br>batch | 25 | 16 | 10 | 20 | 1.078 | 1.59 | 0.0007 |
| cadA+mttA+mfsA<sub>GAP</sub> | Bioreact<br>or,<br>YNB+glyc<br>erol<br>batch | 30 | 16 | 10 | 20 | 2.692 | 0.90 | No growth |
| MC-CBB-IA_<br>cadA+mttA+mfsA<sub>GAP</sub> | Bioreact<br>or,<br>YNB+glyc<br>erol<br>batch | 25 | 16 | 10 | 20 | 6.738 | 3.22 | 0.002 |
| MC-CBB-IA_<br>cadA+mttA+mfsA<sub>GAP</sub> | Bioreact<br>or, YPG<br>batch | 25 | 16 | 10 | 20 | 11.836±0.262 | 3.94±0.006 | 0.004±0.0002 |
| MC-CBB_cadA+mttA+mf<br>sA<sub>GAP</sub> | Bioreact<br>or, YPG<br>batch | 25 | 16 | 10 | 20 | 5.141±0.057 | 1.93±0.072 | 0.003±0.0001 |

### Lower temperature and higher CO_2_ concentration lead to higher itaconic acid production

Standard shake flask experiments have hitherto been conducted at 30°C (with 5% CO_2_), as this is the optimum growth temperature of *K. phaffii* on glucose or methanol. In addition, it has been established that the enzymatic activity of CadA increases with elevated temperatures.^23,24^ However, it is important to note that CO_2_ is a gaseous substrate, and its solubility in liquids is higher at lower temperatures. Furthermore, the specificity of RuBisCO towards CO_2_ increases when temperature is decreased. ^25^ On the other hand, as we demonstrated in our previous study, elevated CO_2_ concentrations boost the itaconic acid production.^16^ Consequently, the performance of the cadA+mttA strain was examined at 25°C and compared to 30°C in shake flask experiments at 10% CO_2_ (Figure 3).

**Figure 3.**
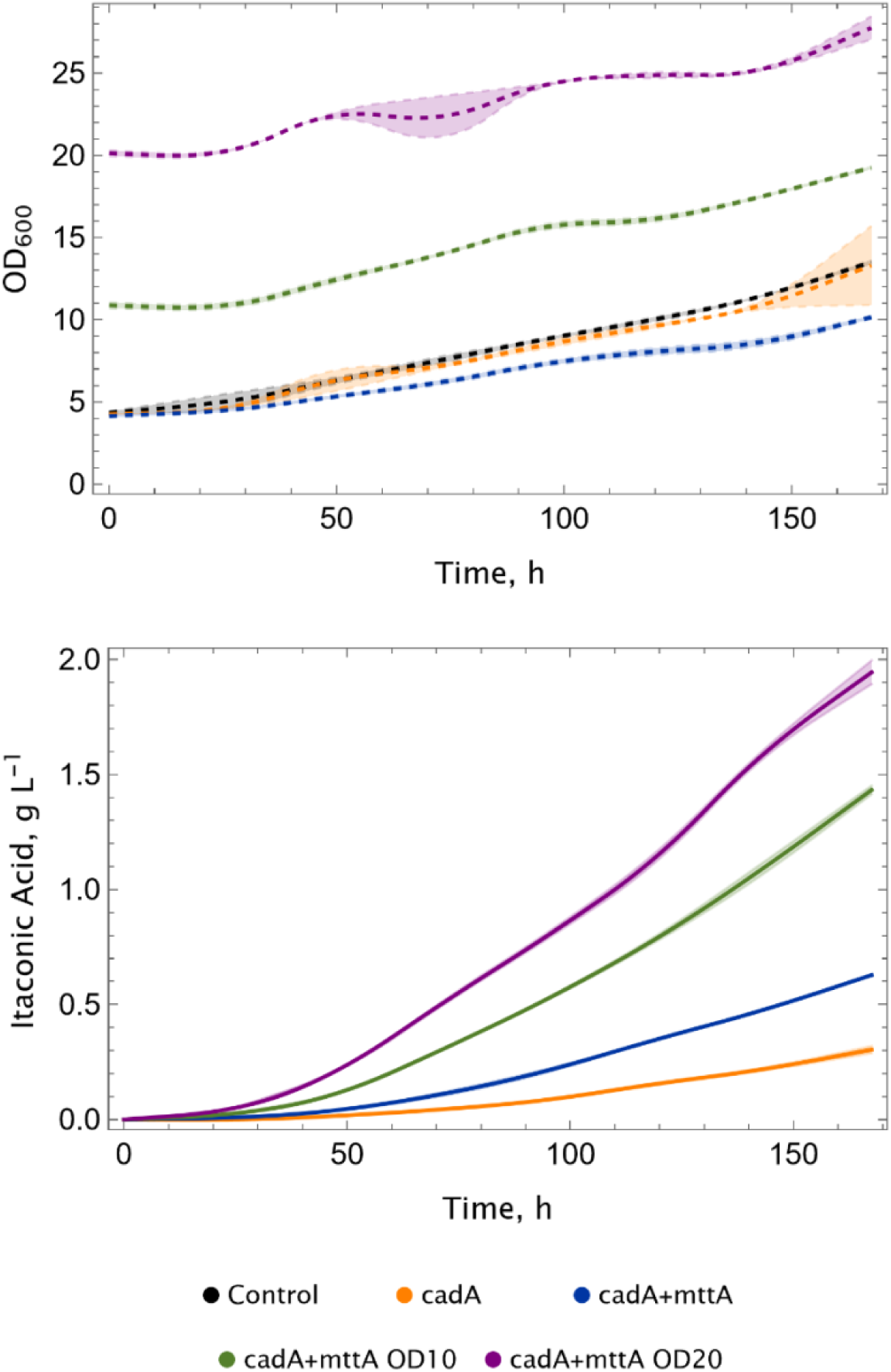
Decreasing temperature improves itaconic acid production. a) Growth and b) itaconic acid production profiles of the cadA+mttA strain. Experiments were conducted in duplicates, CO_2_-shakers supplied with 10% CO_2_. The temperature of the shaker was kept between 25-28°C during the cultivation. Standard deviations (±) are calculated and shown in shades.

A 5°C decrease in temperature resulted in a two-fold impact: Firstly, the growth of the control strain and the production strains was enhanced in comparison with 30°C. In our previous study, cultivating cells with higher initial ODs did not result in growth, and the ODs remained almost unchanged during the cultivation process. Conversely, at a lower temperature, a growth rate of 0.002 h^-1^ was observed at initial OD=20 at 25°C, albeit still lower than the growth rate with initial OD=4. Furthermore, at an initial OD of 4, the growth rate was found to be 0.006 h^-1^, which is 1.5-fold higher than that observed at 30°C. Secondly, the final titers of the itaconic acid obtained was enhanced and reached to approximately 1.95 g L^-1^ (compared to 1.71 g L^-1^ at 30°C with 10% CO_2_).^16^

Due to technical limitations during shake flask cultivations, it was not possible to maintain a constant temperature of 25 °C, which fluctuated between 25 and 28 °C. Consequently, we wanted to assess the impact of temperature in a more controlled environment, and conducted bioreactor cultivations. As anticipated, the results demonstrated the positive impact of reduced temperatures on both growth and itaconic acid production performance (Figure 4). The growth was close to zero (µ=0.0004 h^-1^) for the cultivations at 25°C, whereas a decrease in the biomass was observed during the cultivation at 30°C. The final titers and specific productivity were increased 2.1-fold and 1.6-fold, respectively, compared to 30 °C (Table 1). The highest recorded itaconic acid titer was 1.39 g L^-1^ (versus 0.65 g L^-1^ at 30°C), which is the highest concentration recorded among the experiments conducted so far. Furthermore, the expression levels of key genes involved in the CBB cycle and itaconic acid metabolism were analysed (see Supplementary Figure 2). We took samples at three different time points. The expression levels of the measured genes from the CBB cycle (*PRK*, RuBisCO), the TCA cycle (*CIT1*, *ACO1*, *ACO2*), and itaconic acid metabolism (*cadA*, *mttA*) were found to be elevated at 25°C in comparison to 30°C during the 148 hours of bioreactor cultivation, with the exception of the final time point (194 hours of the cultivation).

**Figure 4.**
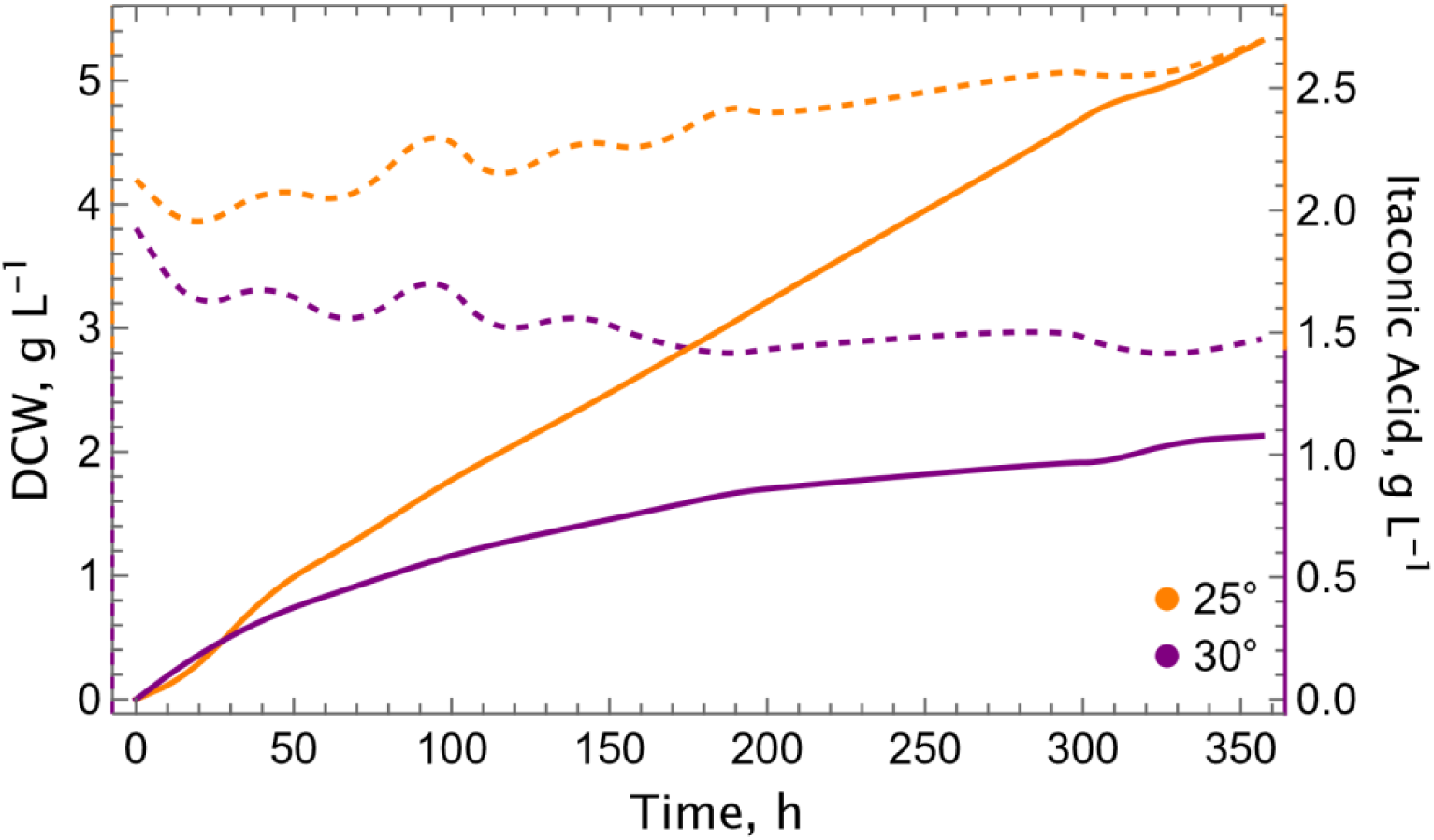
Cultivation at a lower temperature increases itaconic acid production in the bioreactor. Growth (dashed lines) and itaconic acid production (solid lines) profiles at 25°C (green) and 30°C (orange) with the cadA+mttA strain. Experiments were conducted in single runs in lab-scale bioreactors, using 10% CO_2_ in the inlet gas.

### Co-expression of *mfsA* leads to improvement of itaconic acid titers

Following the identification of the key process parameters, we shifted our focus to metabolic engineering to improve the strain efficiency. We introduced the *mfsA* gene to the cad+mttA strain to test whether co-expression would increase the itaconic acid production. Utilising two strong promoters, pFDH1 (methanol inducible) and pGAP (constitutive), the impact of *mfsA* on itaconic acid production in the cadA+mfsA strain was examined in shake flasks.

The co-expression of *mfsA* resulted in a 20% increase in the final itaconic acid titers, yielding approximately 1 g L^-1^ in both the cadA+mttA+mfsA_FDH1_ and cadA+mttA+mfsA_GAP_ strains (Figure 5). Growth was not affected in any of the strains (0.005 h^-1^), specific productivity was similar and varied between 3.1-3.4 mg g^-1^ h^-1^ (Table 1). For further experiments, a representative clone for cadA+mttA+mfsA_GAP_ was selected based on the advantages offered by a constitutive promoter for an exporter.

**Figure 5.**
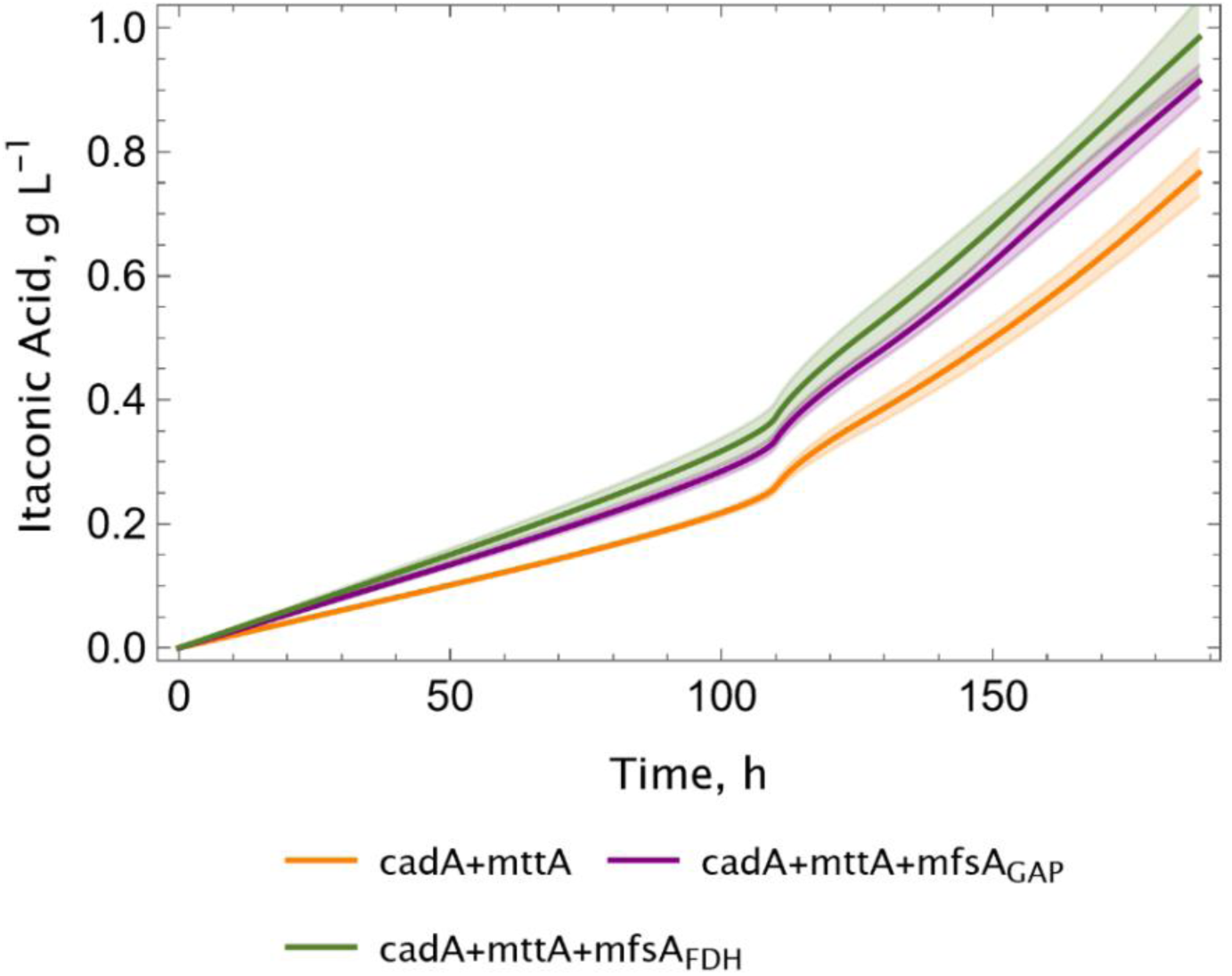
Co-expression of *mfsA* improves the itaconic acid production. Screening was performed in shake flasks at 25-28°C, 5% CO_2_. Seven biological replicates for each construct were screened and standard deviations (±) were calculated and shown in shades.

After verifying the superior performance of the strain with co-expression of *mfsA*, we tested its performance in bioreactor cultivations. We included a bioreactor run at 30°C to compare with 25 °C, thereby investigating the potential benefits of elevated temperatures on itaconic acid production and export (Figure 6). This has been reported recently by Severinsen et al.^24^ where 30-32 °C were reported as optimum temperatures for itaconic acid production by *K. phaffii* on methanol. However, in our experimental setting, 25 °C was found to be more conducive than 30 °C, with the itaconic acid titer reaching 2.70 g L^-1^ after a 360-hour cultivation period (Figure 6). In alignment with our previous bioreactor cultivations conducted at 30°C, the cadA+mttA+mfsA_GAP_ strain did not demonstrate any signs of growth. However, the specific growth rate of the same strain was 0.0007 h^-1^ at 25°C, which is 1.75-fold higher than the strain cadA+mttA (Figure 4), while the specific productivity was 1.2-fold higher (1.59 vs. 1.32 mg g^-1^ h^-1^).

**Figure 6.**
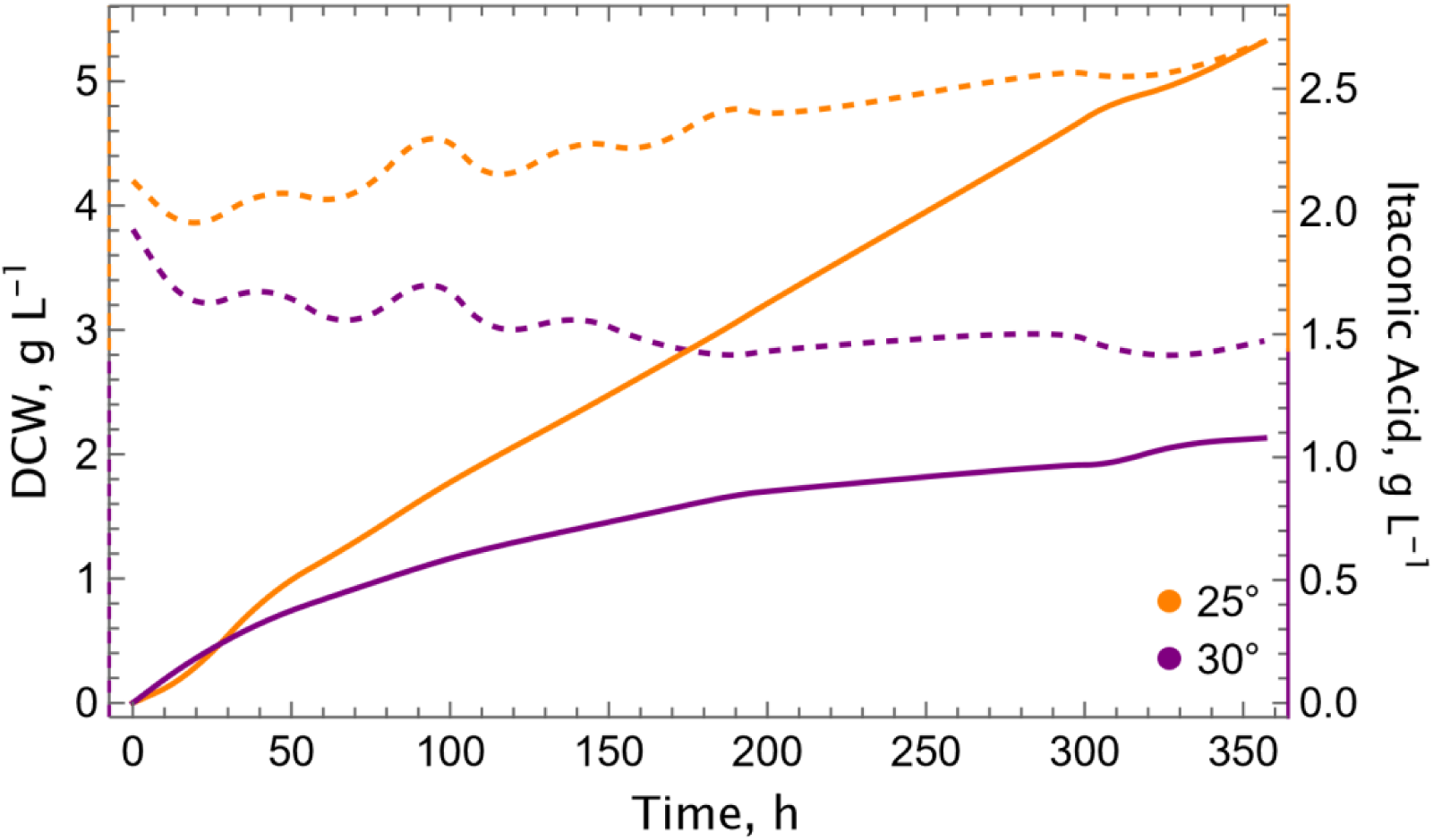
Elevated CO_2_ concentration and decreased temperature improves the tters of itaconic acid production with the cadA+mttA+mfsA_GAP_ strain in bioreactor cultivation. Growth (dashed lines) and itaconic acid production (solid lines) profiles at 25°C and 30°C are shown. Experiments were conducted in single runs in lab-scale bioreactors, using 10% CO_2_ in the inlet gas.

### Improving efficiency of the CBB cycle is beneficial for itaconic acid production

Itaconic acid is a by-product of the TCA cycle, and its production is positively correlated with growth. As was demonstrated above, lowering the temperature can enhance growth. Therefore, we hypothesized that a strain capable of enhanced growth might increase productivity. To test the hypothesis, we constructed a production strain, MC-CBB_cadA+mttA+mfsA_GAP_, using a parental strain that carries multiple copies of RuBisCO and chaperones.^26^ This strain showed superior growth compared to the single-copy strain, that we have employed to construct production strains until this point. The performance of MC-CBB_cadA+mttA+mfsA_GAP_, in addition to the multicopy control strain, was evaluated in shake flask experiments (Supplementary Figure 3).

As anticipated, the MC-CBB_cadA+mttA+mfsA_GAP_ strain exhibited a titer of 1.37 g L^-1^ itaconic acid at 195 hours (equivalent to 2.02 g L^-1^ at 262 hours) during the cultivation period, while the cadA+mttA+mfsA_GAP_ strain reached to a titer of 1.11 g L^-1^ (1.57 g L^-1^ at 262 hours). The productivity of the MC-CBB_cadA+mttA+mfsA_GAP_ strain was found to be 4.15 mg g^-1^ h^-1^, which is a 1.3-fold increase in comparison with the cadA+mttA+mfsA_GAP_ strain (Table 1).

### Increasing the copy number of itaconic acid metabolism improves the titers

After demonstrating that the MC-CBB_cadA+mttA+mfsA_GAP_ strain has a superior performance in the production of itaconic acid, we decided to increase the gene copy number of the itaconic acid metabolism to test whether multiple copies could improve the titers further. Here, either we integrated multiple copies of each gene individually (*cadA*, *mttA* or *mfsA*) or combinations of these genes to construct MC-CBB-IA_cadA+mttA+mfsA_GAP_ strains. We selected random colonies from each transformation and screened them for the final itaconic titers and respective product yields (Figure 7). The integration of *mttA* alone resulted in a substantial growth impairment. This finding is consistent with our previous results, where *mttA* was co-expressed with a strong promoter, resulting in impaired growth.^16^

**Figure 7.**
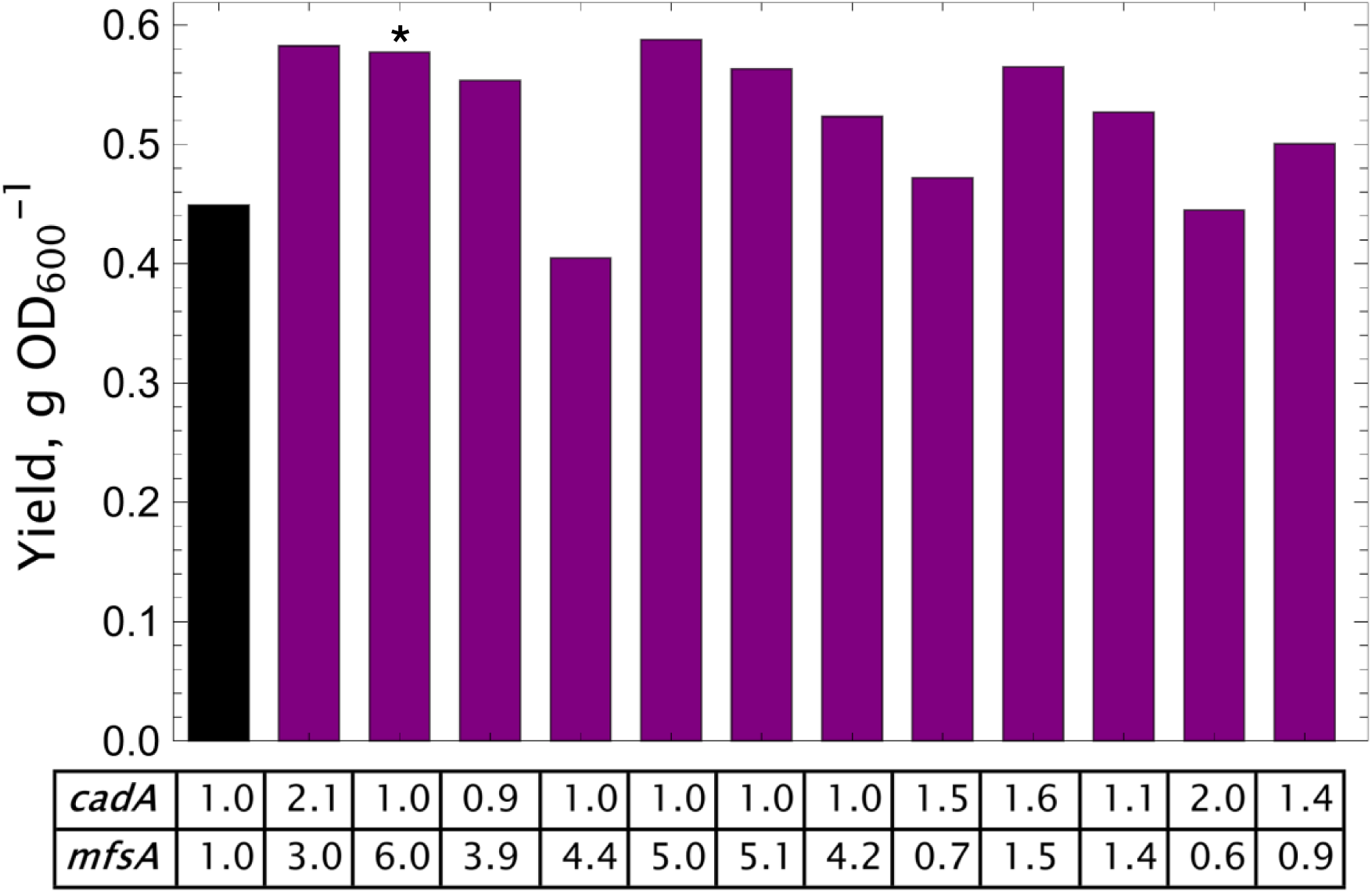
Increasing the copy number of *mfsA* enhances the itaconic acid yield. At least 3 clones from each construct were screened. Screening was performed in shake flasks at 25°C, 5% CO2. The numbers in the table represent the gene copy numbers of the respective genes. Black: control, purple: multicopy clones. Selected clone for further experiments is indicated with an asteriks.

The co-expression of *cadA* alone did not result in a significant enhancement of production and the best results were achieved when multiple copies of *mfsA* were present. This is corroborated by gene copy number (GCN) analysis. The enhanced levels of itaconic acid titers and product yields (g OD_600_^-1^) were consistent with the elevated copies of *mfsA*, exhibiting a specific productivity of 5.19 mg g^-1^ h^-1^ (Table 1). Consequently, the clone from the MC-CBB-IA_cadA+mttA+mfsA_GAP_ strain, which contains around six copies of *mfsA*, was selected for further bioreactor experiments.

### Identifying the bottlenecks in bioreactor cultivations boosts the itaconic acid production

Following the construction of the MC-CBB-IA_cadA+mttA+mfsA_GAP_ strain, bioreactor cultivations were performed utilising the parameters that had been determined in the preceding experiments: 25°C, 10% CO_2_ and 16% dissolved oxygen concentration. However, during bioreactor cultivations, it was still not possible to achieve or exceed the titers and productivity levels obtained in shake flasks. Despite the fact that temperature and dissolved oxygen levels were identified as two pivotal parameters, these efforts were unsuccessful, thus pointing out a limitation.

Our standard bioreactor cultivations included a glycerol batch phase with yeast nitrogen base (YNB) containing 8 g L^-1^ glycerol. In this setup, cells were grown to a DCW of approximately 4 g L^-1^ (OD 20) and the autotrophic growth conditions for the itaconic acid production were initiated with 10% CO_2_. This constitutes a fundamental difference between shake flask and bioreactor cultivations, wherein a preculture in YPG is employed in the shake flasks, as opposed to the YNB + Glycerol batch phase in the bioreactor. In order to address the aforementioned problem, we sought to mimic the conditions of the shake flask screenings in a lab-scale bioreactor cultivation as closely as possible: we employed another approach, where a “batch phase”was performed in shake flasks in YPG instead of YNB+ Glycerol, similar to shake flask cultivations. After growing the cells in YPG, we washed them twice to remove the residuals and inoculated the bioreactor with an OD of approximately 20, and the autotrophic production phase with 10% CO_2_ was initiated.

The analysis of the growth profile revealed the beneficial effect of incorporating a “batch phase”in a complex medium, as it enabled the cells to reach higher biomass titers with a growth rate of 0.004 h^-1^ compared to 0.002 h^-1^ (Figure 8, Table 1). The final titers achieved under these conditions were the highest among the tested conditions and measured as 11.84 g L^-1^. This is consistent with the observed continuous growth, as itaconic acid is a product of the primary metabolism. The results demonstrate a clear enhancement in itaconic acid production for the MC-CBB-IA_cadA+mttA+mfsA_GAP_ strain, with a productivity of 3.94 mg g^-1^ h^-1^. This is a 2-fold increase compared to the control strain (MC-CBB_cadA+mttA+mfsA_GAP_) in YPG batch culture and 1.2-fold higher than the same strain grown in a YNB+Glycerol batch (Table 1).

**Figure 8.**
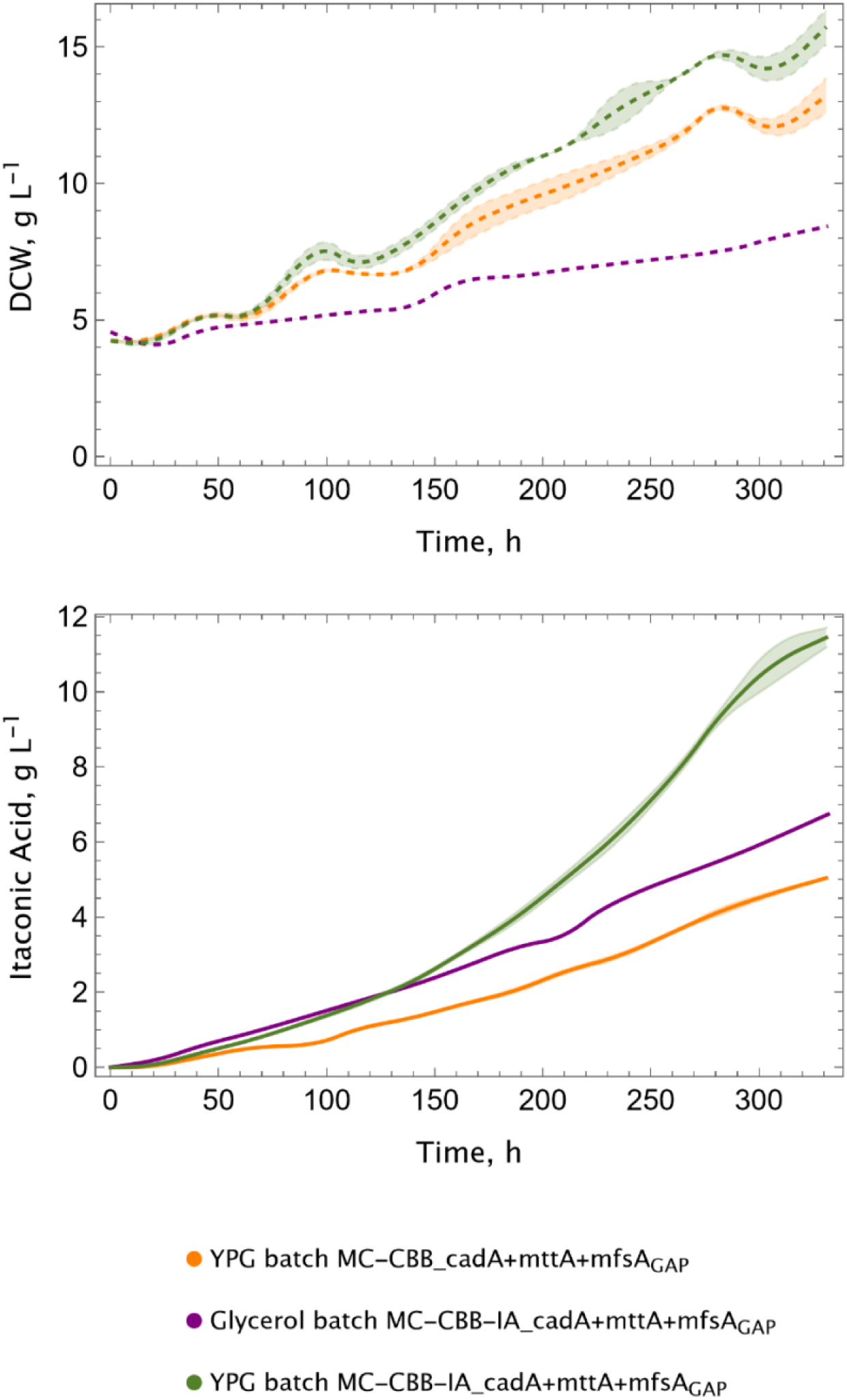
Preculture medium affects the performance of the itaconic acid production strains. Growth (dashed lines) and itaconic acid production (solid lines) profiles. Experiments were conducted at 25°C in duplicates in lab-scale bioreactors, using 10% CO_2_ in the inlet gas.

## Discussion

In this study, we integrate metabolic engineering strategies to optimize flux distribution in heterologous pathways with further bioprocess design focusing on the key process parameters, *i.e.* DO and temperature, to enhance itaconic acid production via CO₂ conversion in bioreactor cultivation.

Previously, we demonstrated that a DO concentration of 8% enhanced itaconic acid production when compared to levels of 20%.^16^ Therefore, here, the investigation was expanded to encompass DO values below 20%, and we included the DO concentrations of 4%, 8%, and 16%, to examine the relationship between oxygen levels and itaconic acid production. The impact of oxygen should be considered at two levels, namely the oxygenation side reaction of RuBisCO and the methanol oxidation involved in the dissimilation. Increasing oxygen availability may potentially decrease the reaction rates of RubisCO on CO_2_ as oxygen serves as an additional substrate, therefore lower oxygen concentrations could be advantageous. However, the cells require oxygen for the methanol oxidation and *K. phaffii* is unable to grow anaerobically. Consequently, a reduction in oxygen availability may lead to a decrease in reducing equivalents, which are produced during the methanol dissimilation process. In this context, a DO level of 16% was found to be the optimum oxygen level with a 1.4-fold increase in itaconic acid production titer.

The second key parameter is the temperature, which had a substantial impact both on production and growth, when it was switched from 30 °C to 25 °C. Previously, we evaluated different initial biomass values to see their effect on itaconic production at 30°C. ^16^ In this setup, even though the production was improved, there was almost no growth; especially when the initial biomass was higher. Conversely, at 25 °C, cells exhibited growth, suggesting a more efficient CBB cycle compared to 30 °C. Furthermore, we examined the impact of temperature on bioreactor cultivations, where the itaconic acid titer reached 1.39 g L^-1^ at 25 °C, representing a 2.1-fold increase compared to 30 °C. The benefit of lower temperature was also reflected in the gene expression levels of the CBB cycle, as at lower temperatures, RuBisCO and PRK were more highly expressed compared to 30 °C. One evident reason for obtaining higher titers and better growth at lower temperature could be the higher gas solubility in the aqueous media. The availability of CO_2_ might have been increased due to the lower temperature, which then leads to an increased carbon flux in the CBB cycle with. A secondary rationale pertains to the catalytic properties of RuBisCO, as shown by Yamori et al. (2006).^27^ They demonstrated that as temperature rises, the carboxylation rate and specificity factor (S_c/o_) decrease. This phenomenon further elucidates why lower temperatures yield superior titers and promote growth. In our previous study, we demonstrated that a balanced co-expression of *mttA*, a mitochondrial membrane transporter, is essential for enhanced production and growth. Here, we further demonstrate the role of another transporter, *mfsA*, leading to higher productivity, confirming previous findings.^24,28^ The enhanced titers observed in the *mfsA*-expressing strain are probably indicative of a bottleneck in the secretion of itaconic acid, rather than a deficiency in its production. In fact, increasing the copy number of *mfsA* led to a substantial enhancement in itaconic acid production, thereby enhancing the pull effect. Conversely, the introduction of additional copies of cadA or mttA did not result in a significant change in itaconic acid levels, whereas higher copy numbers of mttA even led to a substantial growth impairment. This outcome is consistent with prior findings.^16,24^ The enhanced expression of mttA is likely to cause a depletion of *cis*-aconitate, an intermediate in the TCA cycle, leading to diminished TCA cycle activity. Consequently, it is essential to balance the expression of heterologous genes, particularly when they interfere with native essential metabolic pathways, to ensure efficient product synthesis.

The results of integrating multiple copies of the CBB cycle and itaconic acid metabolism genes, showed that it is crucial to optimize the balance of carbon flux from the CBB cycle to biomass and itaconic acid production precursors. Increasing the copy numbers of the pivotal gene, RuBisCO, and its chaperones, GroEL and GroES, not only improved the growth but also led to a substantial increase in the production performance of the synthetic autotrophic strain. In a previous study, adaptive evolution led to mutations in PRK with reduced expression levels and enzyme activity, and reverse engineering that in the initial CBB cycle strain confirmed increased growth relative to the parent strain.^29^ However, it was observed that this strain did not exhibit superiority in terms of itaconic acid production, despite its enhanced growth capacity.^16^ To investigate whether increasing the copies of *mfsA* would lead to a further improvement with a pull effect, the previously mentioned reverse engineered strain was used to integrate itaconic acid metabolism with additional copies of *mfsA*. While higher titers were achieved due to faster growth, the specific productivity of the reverse engineered strain was lower than that of the parent strain, (MC_IA+cadA_mttA_mfsA_GAP_) (Supplementary Table 1). These findings indicate that increasing the growth rate does not necessarily increase the productivity and that a balanced carbon flux between growth and production is essential. The significance of a balanced metabolism is further underscored by the integration of multiple copies of *cadA*, *mttA*, or *mfsA* from the itaconic acid metabolism. The integration of *mfsA*, accompanied by *cadA*, exhibited the highest productivity and titer, while the introduction of additional copies of *cadA* alone did not enhance the strain’s performance.

Finally, we calculated the key process parameters of the bioreactor cultivation, where an itaconic acid titer of approximately 12 g L^-1^ was reached with the highest productivity among the tested conditions in bioreactors (3.94 mg g^-1^ h^-1^) (Table 1). Total product yield on methanol, that was used for the NADH production via the dissimilation, was 0.09 g g^-1^ while the net CO_2_ production was 0.25 mmol CO_2_ g^-1^ h^-1^. Overall, 52 g MeOH was utilized during the cultivation, while 34 g CO_2_, 11 g biomass and 5 g itaconic acid was produced, showing the bioprocess is still CO_2_-positive. However, in the context of a process in which methanol is produced via the electrochemical reduction of CO_2_, there is considerable potential for itaconic acid production with a synthetic autotrophic yeast strain to become a CO_2_-negative process. Additionally, engineering the methanol dissimilation pathway by replacing alcohol oxidase with an alcohol dehydrogenase can be an alternative solution. This modification would lead to the production of an extra mole of NADH, thereby reducing the methanol requirement and contributing to the achievement of CO_2_-neutral or negative bioprocesses.^30^

## Conclusion

In this study, we demonstrated that a synthetic autotrophic *K. phaffii* is able to produce itaconic acid by the direct conversion of CO_2_, achieving approximately final 12 g L^-1^ titers in the bioreactor cultivations. We showed that balancing the host metabolism considering the interplay of the heterologous metabolism with the native metabolic pathways is essential to reach higher production performance. Additionally, beyond metabolic engineering, precise bioprocess design was one of the key steps in enhancing productivity.

The current status of the CO_2_-based process is not yet able to compete with the traditional fermentation processes, where first- or second-generation feedstocks are used. However, we believe our results highlight the potential of single-carbon substrate-based processes, utilizing CO₂ directly or in combination with other CO₂-derived substrates like methanol or formate, to develop CO₂-neutral or negative bioprocesses in line with future sustainability goals.

## Methods

### Construction of the strains

The synthetic autotrophic *K. phaffii* strain described by Gassler et al. (2020)^22^ was used as the host to construct the organic acid producing strains. The construction of the strain carrying *cadA* and *mttA* was described by Baumschabl et al. (2022).^16^ The *mfsA* gene encoding a cytoplasmic itaconic acid exporter was inserted into the *GUT1* locus by CRISPR-Cas9^31^ in the cadA+mttA strain. Two different promoters (pFDH1 and pGAP) were tested to find the optimal expression of *mfsA*. Plasmid constructions and transformations were performed by Golden Gate Assembly (GGA) and the CRISPR-Cas9 system described in Gassler et al. (2019).^29^ Integration to the correct loci was verified by PCR.

Multiple copies of *cadA, mttA, mfsA*, along with combinations of these genes and RuBisCO and its chaperones were performed as described by Severinsen et al. (2024). ^24^ A list of the strains used in this study is given in Table 2.

**Table 2.** *K. phaffii* strains used in this study.

| Strain | Genotype | References |
| --- | --- | --- |
| Control | CBS7435 ΔAOX1::pAOX1_TDH3 + pFDH1_PRK + pALD4_PGK1 | Gassler et al 2020<sup>22</sup> |
| | ΔDAS1::pDAS1_RuBisCO +<br>pPDC1_groEL + pPP1B_groES<br>ΔDAS2::pDAS2_TLK1 + pRPS2_TPI1 | |
| cadA | Control + pAOX1_cadA | Baumschabl et al<br>2022<sup>16</sup> |
| cadA+mttA | Control +<br>pAOX1_cadA+pPOR1_mttA | Baumschabl et al<br>2022<sup>16</sup> |
| cadA+mttA+mfsA<sub>FDH1</sub> | Control + pAOX1_cadA +<br>pPOR1_mttA + pFDH1_mfsA | This study |
| cadA+mttA+mfsA<sub>GAP</sub> | Control + pAOX1_cadA +<br>pPOR1_mttA + pGAP_mfsA | This study |
| MC-CBB_<br>cadA+mttA+mfsA<sub>GAP</sub> | Control + Multicopy CBB<sup>26</sup> +<br>pAOX1_cadA + pPOR1_mttA +<br>pGAP_mfsA | This study |
| MC-IA_<br>cadA+mttA+mfsA<sub>GAP</sub> | Control + Multicopy itaconic<br>acid+pAOX1_cadA + pPOR1_mttA +<br>pGAP_mfsA | This study |
| MC-CBB-IA_<br>cadA+mttA+mfsA<sub>GAP</sub> | Control + Multicopy CBB<sup>26</sup> - itaconic<br>acid+pAOX1_cadA + pPOR1_mttA +<br>pGAP_mfsA | This study |
| RE_cadA+mttA | Reverse engineered control<sup>29</sup> +<br>pAOX1_cadA + pPOR1_mttA + | This study |
| RE_MC_IA_cadA+mttA+mfsA | Reverse engineered control<sup>29</sup> +<br>Multicopy itaconic<br>acid+pAOX1_cadA + pPOR1_mttA +<br>pGAP_mfsA | This study |

### Shake flask cultivations

Shake flask cultivations were carried out in 100 mL narrow-neck flasks with cotton caps at 25 or 30°C with 5 or 10% constant CO_2_ supply, shaken at 180 rpm in 22 mL buffered YNB (3.4 g L^-1^, pH 6) supplemented with 10 g L^-1^ (NH_4_)_2_SO_4_ as the nitrogen source. Cells were inoculated with the target starting OD_600_ (4 or 20) and induced with 0.5% (v/v) methanol at the start of the main culture, and adjusted to 1% (v/v) methanol after the first sampling until the end of the cultivations. Cell growth (OD_600_) was monitored during the cultivation and extracellular metabolite concentrations (methanol and itaconic acid) were measured by high-performance liquid chromatography (HPLC) and culture volume was corrected for evaporation by the addition of water.

### Bioreactor cultivations

Bioreactor cultivations were conducted in 1.4 L DASGIP reactors (Eppendorf). pH was kept at 6.0 by using 2 mol L^-1^ NaOH or 5 mol L^-1^ KOH. Dissolved oxygen (DO) concentration was controlled by adjusting the inlet oxygen concentration, stirrer speed and inlet gas flow whereby 5%, 200 rpm and 6 sL h^-1^ were the minimal setpoints, respectively. DO was set to 4, 8, or 16% to explore the effect of oxygen concentration on growth and itaconic acid production. The autotrophic cultivation was performed from beginning on using 10% CO_2_ in the inlet gas and 0.5 % methanol. After the first sample (appr. 16 h) methanol concentration was adjusted to 1%. Cultivations were conducted using YNB media supplemented with 10 g L^-1^ (NH_4_)_2_SO_4_ as the nitrogen source and buffered using 100 mmol L^-1^ phosphate buffer at pH 6. Temperature was set either to 25 or 30°C.

The bioreactor cultivations were performed in two different experimental setups: 1) A YPG preculture was inoculated to an OD of 1 in YNB media including 8 g L^-1^ glycerol. The batch phase was performed to reach an end biomass of approximately OD_600_ of 20 (ca 4 g L^-1^), based on a yield on glycerol of approximately 0.5. Following the batch end, cultures were induced with 0.5 % (v/v) methanol and provided with constant supply of 10% CO_2_ in the inlet gas starting the autotrophic cultivation. 2) A YPG preculture was inoculated to an OD_600_ of 20 and the autotrophic process conditions were directly initiated, skipping the batch phase on YNB with glycerol.

In both scenarios, methanol concentration was adjusted to 1% at the first sample points (after appr. 24 h). From this time on samples were taken daily including OD, dry cell weight (DCW), and HPLC samples. After each sampling the methanol concentration was adjusted to 1% (v/v). Overall specific growth rate and production rates in the shake flasks and bioreactor cultivations were calculated according to Equation 1 and 2:

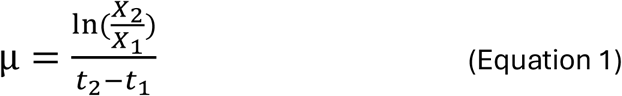

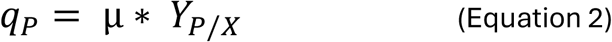

For the nongrowing strains, production rates were calculated according to Equation 3:

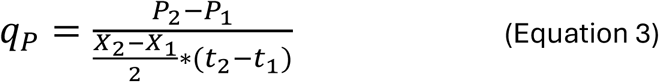

µ is the specific growth rate (h^-1^), X is the cell concentration (g L^-1^) at t (h) of the cultivation, P is the product concentration (mg L^−1^) at t (h) of the cultivation, q_P_ is the specific production rate (mg g^-1^ h^-1^), Y_P/X_ is the product yield (mg g^-1^ h^-1^).

### HPLC measurements

HPLC measurements were performed by using a a Biorad Aminex HPX-87H HPLC column (300 x 7.8 mm) (Baumschabl et al 2022). Samples were centrifuged at 4000 g for 10 min, and the supernatant of each sample was mixed with 40 mM H_2_SO_4_ resulting in a final concentration of 4 mM. Samples were vortexed and centrifuged at full speed (16100 g) for 5 min at room temperature. Following the centrifugation, they were filtered using a 0.22 µm filter into the vials for the HPLC analysis.

### Genomic DNA extraction and gene copy number analysis

Genomic DNA (gDNA) of the *K. phaffii* strains was extracted from overnight cultures using the Wizard Genomic DNA purification kit (Promega Corp., USA) according to the manufacturer’s instructions. The quality, purity and concentration of the isolated gDNA was verified with Nanodrop.

Following the genomic DNA extraction, an RT-PCR was carried out using 2x qPCR S’Green BlueMix (Biozym Blue S’Green qPCR Kit). gDNA was mixed (2.7 ng µL^-1^ in 3 µL) with respective primers (*cadA, mttA, mfsA*; 0.4 µL from 10 mM stocks), 2x qPCR S’Green BlueMix (5 µL) and water (up to 10 µL). Primer sequences are given in Supplementary Table 1. All samples were analyzed in triplicates with non-template controls. The copy numbers of the respective genes were estimated in comparison to the copy number of the parent strain using the ΔΔCT method^32^.

## Acknowledgements

The COMET center acib: Next Generation Bioproduction is funded by BMIMI, BMWET, SFG, Standortagentur Tirol, Government of Lower Austria and Vienna Business Agency in the framework of COMET – Competence Centers for Excellent Technologies. The COMET Funding Program is managed by the Austrian Research Promotion Agency FFG. We thank the Austrian Science Fund for support to DM and MB (Grant-DOI 10.55776/W1224, Doctoral Program on Biomolecular Technology of Proteins (BioToP)), as well as support to ÖA (Grant-DOI 10.55776/M2891).

## Author contributions

ÖA designed and performed the construction of strains and their evaluation. MB performed the construction of strains. LL supported cloning, cultivation and analysis of strains. DM conceived of the study and supervised the project with ÖA. ÖA and DM wrote the manuscript with support by all authors. All authors read and approved the final manuscript.

