## Supplementary File for "Synthetic autotrophic yeast enables high itaconic acid production from CO_2_ via integrated pathway and process design"

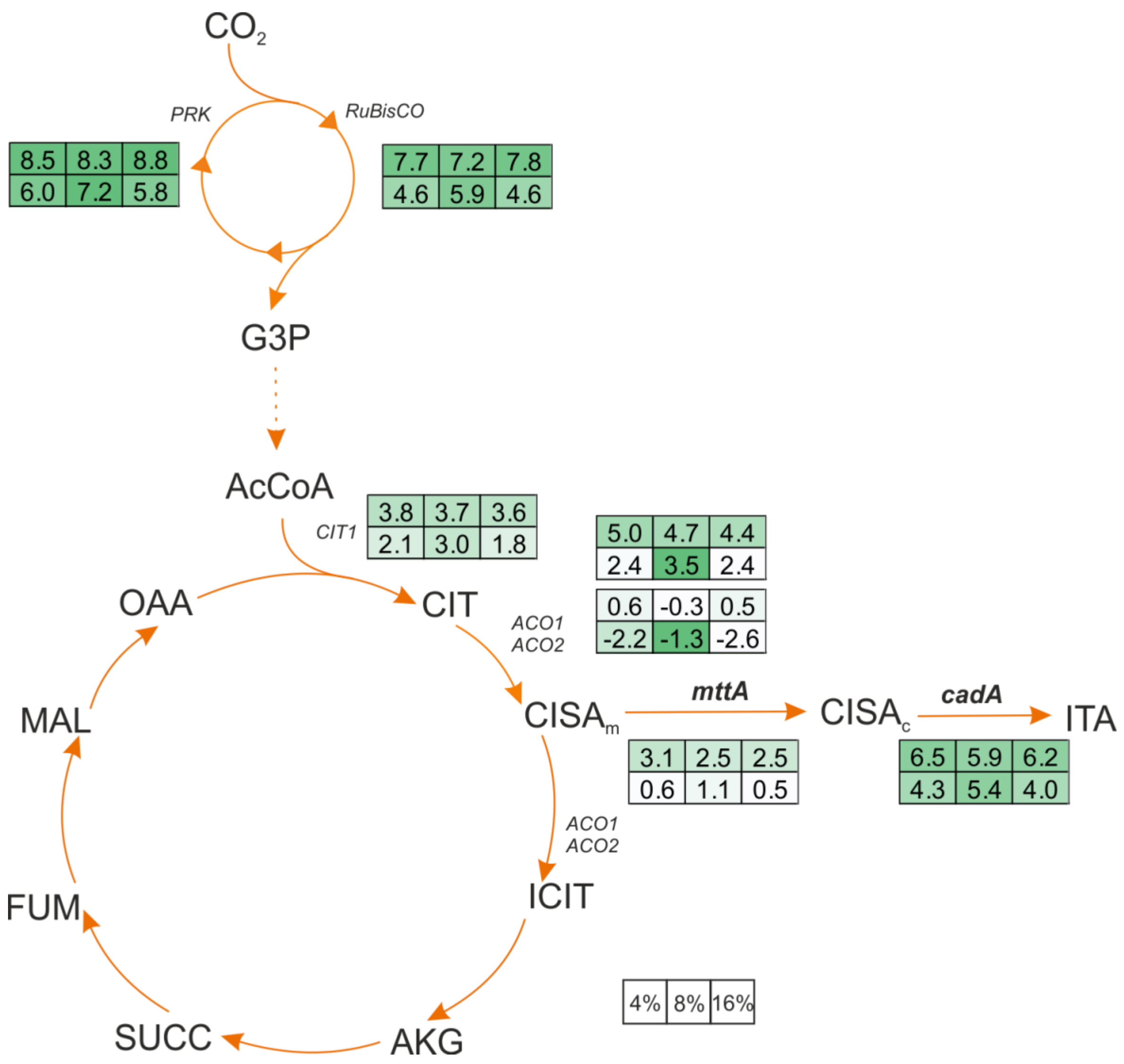


**Supplementary Figure 1**. Gene expression levels of the cadA+mttA strain at different oxygenation conditions at 30°C in the bioreactor cultivation using 5% $CO_2$ in the inlet gas stream. log2-fold change values in the gene expression levels are given. All expression levels are normalized against the strain's own actin (*ACT1*) expression. First and second row values belong to 103 h and 198 h samples, respectively. First, second, and third column values belong to the cultivations performed at 4%, 8% and 16% dissolved oxygen concentration, respectively. AcCoA: acetyl-coenzyme A, AKG: alpha-ketoglutarate, *cadA*: cis-aconitate decarboxylase, $CISA_c$: cytosolic cis-aconitate, $CISA_m$: mitochondrial cis-aconitate, CIT: citrate, ICIT: isocitrate, FUM: fumarate, ITA: itaconic acid, MAL: malate, *mttA*: mitochondrial tricarboxylic acid transporter, OAA: oxaloacetate, *PRK*: phosphoribulokinase, *RuBisCO*: ribulose 1,5-bisphosphate carboxylase/oxygenase, SUCC: succinate, peroxisome. c: cytosol, m: mitochondria.

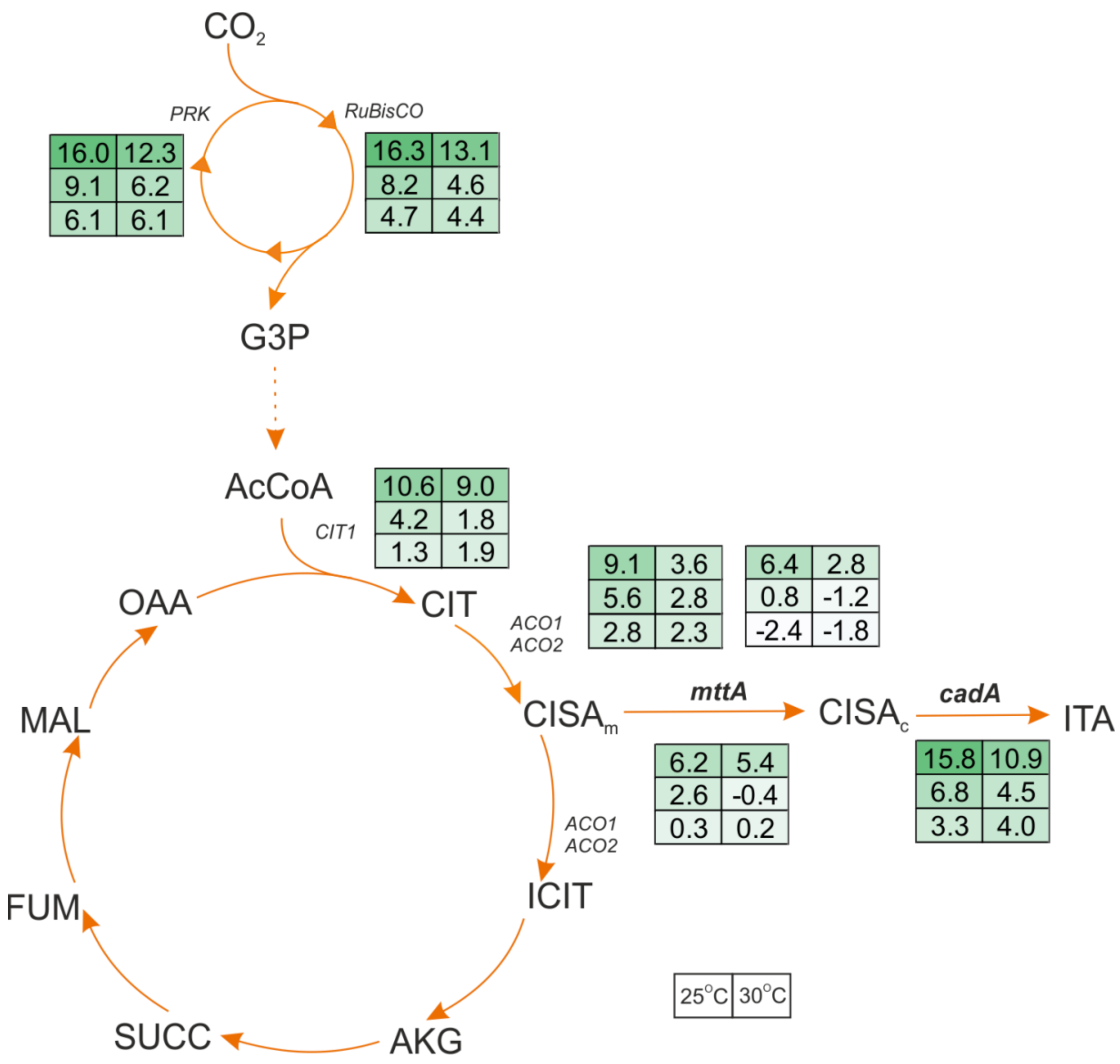


**Supplementary Figure 2**. Gene expression levels at different temperatures of the cadA+mttA strain at 25°C and 30°C in the bioreactor cultivation using 10% $CO_2$ in the inlet gas stream. log2-fold change values in the gene expression levels are given. All expression levels are normalized against the strain's own actin (*ACT1*) expression. First, second and third row values belong to 27 h, 148 h, and 194 h samples, respectively. Left and right column values belong to the cultivations at 25 and 30°C, respectively. AcCoA: acetyl-coenzyme A, AKG: alpha-ketoglutarate, *cadA*: cis-aconitate decarboxylase, $CISA_c$: cytosolic cis-aconitate, $CISA_m$: mitochondrial cis-aconitate, CIT: citrate, ICIT: isocitrate, FUM: fumarate, ITA: itaconic acid, MAL: malate, *mttA*: mitochondrial tricarboxylic acid transporter, OAA: oxaloacetate, *PRK*: phosphoribulokinase, *RuBisCO*: ribulose 1,5-bisphosphate carboxylase/oxygenase, SUCC: succinate, peroxisome. c: cytosol, m: mitochondria.

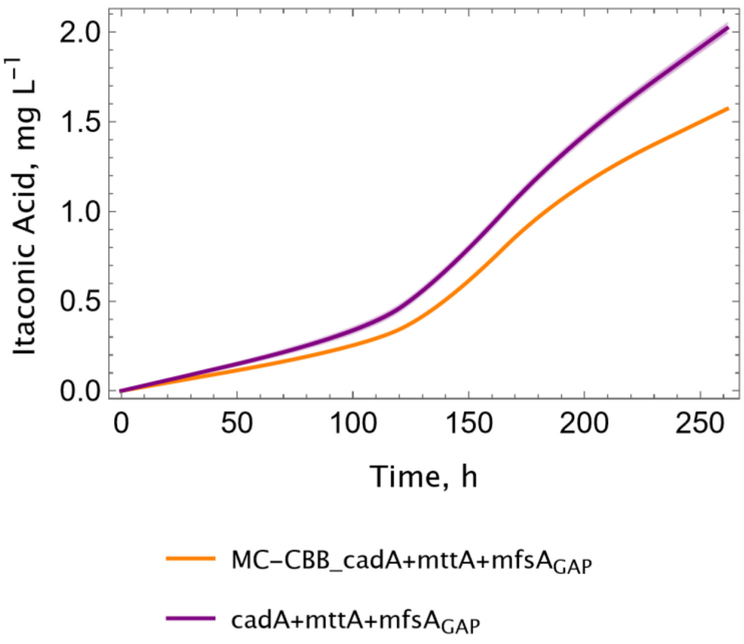


**Supplementary Figure 3. Higher copy number of the CBB cycle improves itaconic acid production**. Itaconic acid production profile of the cadA+mttA+mfsA$_{GAP}$ and MC-CBB_cadA+mttA+mfsA$_{GAP}$ strains. Screening was performed at shake flask at 25°C, with 5% $CO_2$. Four biological replicates for each construct were screened and standard deviations (±) were given in shades.

**Supplementary Table 1.** Growth and itaconic acid characteristics of the reverse engineered[1] (RE) strain with multicopy of *mfsA*. Data is calculated using the screening data at 25°C, with 10% $CO_2$. Genotypes of the strains are given in Table 2.

| Strain | Final Titer (g $L^{-1}$) | Productivity, $q_p$ (mg g $DCW^{-1}$ $h^{-1}$) | Growth rate, μ ($h^{-1}$) |
|---|---|---|---|
| RE | - | - | 0.012 |
| RE_cadA+mttA | 0.826 | 1.100 | 0.010 |
| RE_MC_IA_cadA+mttA+mfsA | 1.365±0.324 | 1.784±0.344 | 0.010±0.000 |
| MC-IA_ cadA+mttA+$mfsA_{GAP}$ | 0.870±0.284 | 2.458±0.273 | 0.007±0.001 |
